# Switch from inflammatory signaling to excessive extracellular matrix remodeling in fibrotic fibroblasts attenuates monocyte tissue infiltration

**DOI:** 10.64898/2026.11.17.767293

**Authors:** Sabine de Winter, Elsa Neubert, Erik HJ Danen

## Abstract

Tissue damage induces infiltration of monocytes that differentiate into macrophages and cooperate with fibroblasts in tissue repair. If repair fails and tissue stress persists, a feedforward loop involving high contractility of activated fibroblasts, extracellular matrix (ECM) stiffening, and TGF-β1 signaling can lead to chronic fibrosis characterized by hypertrophic scarring. In advanced fibrotic tissues areas devoid of immune infiltration have been detected but how the interplay between chemotactic signals versus ECM remodeling impacts monocyte infiltration and spatial positioning in such late-stage fibrotic tissues is unresolved. To address this, we designed a series of 3D co-culture formats incorporating fibrillar collagen I-based ECM, human fibroblasts, and primary human monocytes. RNA-seq, secretome analysis, and confocal microscopy showed that collagen-embedded fibroblasts strongly upregulated inflammatory signaling as compared to 2D fibroblast cultures. Incorporation of fibroblasts enhanced collagen fiber network porosity and stimulated monocyte entry into collagen hydrogels and their migration within the collagen network. TGF-β1 activation of collagen-embedded fibroblasts caused a striking switch from inflammatory signaling to an ECM remodeling signature, which was accompanied by local ECM accumulation around fibroblasts and a pronounced increase in collagen fiber network pore size. This did not affect overall fibroblast-stimulated monocyte migration through the ECM, but it prevented monocytes from crossing liquid-hydrogel barriers to penetrate the collagen network. Moreover, it caused excessive ECM remodeling near clusters of fibroblasts, which locally arrested monocyte migration. These findings indicate that excessive fibroblast activation in fibrotic tissues can shift the balance from chemotactic cues toward biomechanical constraints, which limits monocyte infiltration and locally restricts monocyte migration.

## Introduction

Organ fibrosis represents the outcome of a dysregulated wound healing response and is characterized by excessive deposition of extracellular matrix (ECM), leading to disrupted tissue architecture, tissue stiffening, and loss of organ function^1^. Although certain diseases are primarily defined as fibrosis, such as Idiopathic Pulmonary Fibrosis and Systemic Sclerosis, fibrotic pathology typically arises as a secondary complication following persistent injury and inflammation. Sustained pro-fibrotic signaling drives progressive ECM production and tissue remodeling, which can lead to dysfunction of organs including the lung, liver, and kidney^2^. Moreover, dense and stiffened fibrotic stroma in tumors contributes to disease progression and therapy resistance, thereby representing a major challenge for cancer treatment^3^. Consequently, fibrosis contributes substantially to global mortality^4^.

The complex fibrotic tissue microenvironment is composed of the ECM and multiple interacting cell types. Our incomplete understanding of the interactions between these components hampers the development of anti-fibrotic therapies^5^. Fibroblasts play a central role in fibrosis through their capacity to remodel the ECM^6^. ECM remodeling includes the deposition, assembly, degradation, contraction, alignment, and crosslinking of ECM components, thereby modifying matrix composition, architecture, and mechanical properties^7^. Under normal conditions, resident fibroblasts maintain tissue homeostasis. However, in response to cues such as cytokines, growth factors, and increased tissue stiffness, fibroblasts become activated and can differentiate into myofibroblasts^8^. Amongst these, transforming growth factor-beta 1 (TGF-β1) is a central regulator of fibroblast activation and myofibroblast differentiation^9^. Activated fibroblasts are characterized by proliferation, elevated cytokine secretion, and production of ECM proteins such as type I and IV collagen and fibronectin, and the subset of myofibroblasts is additionally characterized by their strong contractility associated with upregulated alpha-smooth muscle actin (*a*-SMA) expression^10,11^. Moreover, fibroblasts can reinforce their activation through autocrine and paracrine signaling, thereby amplifying and sustaining a profibrotic niche^12,13^.

Beyond their role in ECM production and tissue remodeling, fibroblasts are increasingly recognized as immunoregulatory cells that shape the inflammatory microenvironment^14^. Fibroblasts can secrete cytokines, chemokines, and growth factors, which recruit immune cells and regulate their functional state^15^. Fibroblasts also regulate the spatial bioavailability of inflammatory signals by remodeling the ECM and thereby releasing inflammatory signals stored in the ECM^16^. Among the immune populations implicated in fibrosis, monocytes are critical mediators of disease progression^17^. Circulating monocytes are recruited from the bloodstream into inflamed and fibrotic tissues in response to chemotactic gradients. Fibroblasts produce several chemokines and cytokines, including C-C motif chemokine ligand 2 (CCL2), which promote the recruitment of circulating monocytes into inflamed tissue, linking fibroblast activation to immune infiltration^18^. CCL2 signals through C-C chemokine receptor type 2 (CCR2) expressed on monocytes and this interaction has been implicated in fibrosis across multiple organ systems^19–21^. Following tissue entry, monocytes differentiate into macrophages whose phenotypes are shaped by local microenvironmental cues, including soluble factors and tissue mechanics^22,23^. Fibroblast-derived cytokines such as IL-1β, TGF-β1, and IL-10, together with fibroblast-mediated ECM remodeling and matrix stiffening, critically contribute to macrophage polarization and function^24^. In turn, macrophages further modulate fibroblast activation and matrix remodeling by secreting inflammatory and profibrotic mediators^25^. Thus, the reciprocal interplay between fibroblasts, infiltrating monocytes, macrophages, and the remodeled ECM establishes a dynamic feedback loop that drives fibrotic progression.

The mechanisms by which monocytes infiltrate and distribute through fibrotic tissues remain incompletely understood. Previous *in vitro* studies demonstrated that fibroblasts can chemotactically recruit monocytes^26,27^ and that the biomechanical characteristics of the ECM influence monocyte migration^28^. It has been shown that ECM remodeling by individual fibroblasts, including fiber alignment, contraction, and tunnel formation, can mechanically guide monocyte and macrophage migration^29,30^. At the tissue level, however, spatial studies in fibrotic diseases have identified myofibroblast- and ECM-rich regions with relatively low immune-cell infiltration^31–33^, while similar immune-restricted stromal niches in tumors have been associated with high fibroblast density and dense or aligned collagen networks^34–36^. Yet the exact role of activated fibroblasts in immune exclusion is unclear. In particular, it remains unresolved how extensive fibroblast-driven ECM remodeling and altered paracrine signaling in areas densely populated with activated fibroblasts together regulate monocyte infiltration^37–39^

Here, we design three-dimensional (3D) co-culture setups incorporating fibrillar collagen I-based ECM, human fibroblasts, and primary human monocytes to address these issues using RNA-seq, secretome analysis, and confocal microscopy. We find that collagen-embedded fibroblasts adopt an inflammatory phenotype that stimulates monocyte recruitment and migration. TGF-β1 activation causes a switch from inflammatory signaling to ECM remodeling that does not affect monocyte migration per se, but rather prevents monocytes from crossing hydrogel interfaces and extensively remodeled areas in fibroblast-dense regions. These findings indicate that monocyte infiltration in advanced fibrotic tissues may be attenuated by reduced attraction and the introduction of biomechanical barriers that cannot be navigated by monocytes.

## Materials and Methods

### Cell culture

BJ human foreskin fibroblasts were obtained from ATCC (CRL-2522) and cultured in DMEM (11885, Gibco) supplemented with 10% fetal bovine serum (FBS, S-FBS-NL-015, Serana), 1X Glutamax (35050061, Gibco) and 25 U/mL penicillin and streptomycin (15070063, Gibco) at 37°C, 5% CO_2_. Fibroblasts were subcultured twice a week and used between passages 3 and 12. For experiments, fibroblasts were cultured in the absence of TGF-β1 or preincubated for 3 days with 10 ng/mL TGF-β1 (100-21-10UG; Peprotech) and then used in the experiment for an additional 3 days in the presence of 10 ng/mL TGF-β1.

Peripheral blood mononuclear cells (PBMCs) were isolated from buffy coats or leukapheresis products (Leukopacks) obtained from healthy donors (Sanquin, Amsterdam, the Netherlands) using Ficoll density-gradient centrifugation. Buffy coats were diluted 1:1 with Hanks’ Balanced Salt Solution (HBSS; 14175095, Gibco), whereas Leukopacks were diluted 2:1 with wash buffer consisting of Dulbecco’s phosphate-buffered saline (DPBS) supplemented with 2% FBS. Subsequently, 35 mL of diluted buffy coat or Leukopack suspension was carefully layered onto 13 or 15 mL of Ficoll-Paque PLUS (GE17-5442-02, Cytiva), respectively, in 50 mL centrifuge tubes. Samples were centrifuged at 400 × g at room temperature for 25 min (buffy coats) or 30 min (Leukopacks). Following centrifugation, the PBMC-containing interphase was collected, and cells were washed three times with RPMI supplemented with 1% FBS (buffy coats) or DPBS supplemented with 2% FBS (Leukopacks). Viable cells were counted using Trypan blue exclusion (15250061, Gibco).

Monocytes were isolated from freshly prepared PBMCs by positive Magnetic-Activated Cell Sorting (MACS) using CD14 MicroBeads (130-118-906, Miltenyi Biotec) according to the manufacturer’s protocol. PBMCs were washed with MACS buffer (DPBS with 0.5% bovine serum albumin (BSA, A9647, Sigma-Aldrich) and 2 mM EDTA (E9884-100G, Sigma-Aldrich)) and centrifuged at 300 x g for 10 min. Then, cells were resuspended in MACS buffer and processed at 4°C to minimize cell activation. Cells were mixed and incubated with CD14-microbeads for 15 min, washed with MACS buffer, and applied to a MACS LS separation column placed in a QuadroMACS™ magnetic separator (130-091-051, Miltenyi Biotec). The column was washed with MACS buffer to remove unlabeled cells, removed from the magnetic field and retained CD14+ cells were collected. Next, cells were washed with RPMI medium (61870-036, Gibco), and frozen in batches in 90% FBS and 10% DMSO (102952, Sigma-Aldrich). Cell yield and viability were assessed using Trypan blue and aimed for > 90%.

### 3D cell cultures and monocyte infiltration

For hydrogel anchoring, 384-well µ-clear plates (781091, Greiner) were coated with 1 mg/mL dopamine hydrochloride (H8502-5G, Sigma-Aldrich) in 10 mM Tris (pH 8.6) prepared in DPBS for 16 h at room temperature to generate a polydopamine layer. Wells were washed three times with DPBS and dried completely. Fibroblasts or fibroblasts exposed to TGF-β1 (10 ng/mL) for 3 days were suspended (5*10^5^ cells/mL) in a 2 mg/mL neutralized collagen mixture consisting of complete DMEM, rat tail collagen I (354249, Corning), and 10× PBS, and the pH was adjusted to 7.5 using NaOH. 20 µL of the resulting collagen gel/fibroblast suspension was dispensed into each well and polymerized for 20 min at 37°C, 5% CO₂. To reduce cell sedimentation during polymerization, the plate was flipped upside down after 5 min. Following polymerization, hydrogels were washed twice with complete medium and maintained in DMEM with or without TGF-β1 (10 ng/mL) for 24 h at 37°C, 5% CO₂ prior to monocyte seeding.

CD14+ monocytes were thawed in RPMI medium supplemented with 10% FBS and 25 U/mL penicillin and streptomycin, stained with 1 µM fluorescent CellTracker Green (C2925, Invitrogen) for 20 min at 37 °C, washed once with DPBS, and resuspended in complete DMEM at 1×10^6^ cells/mL. For seeding on top of 3D collagen/fibroblast cultures, 30 µL medium was removed from the hydrogels, and 30 µL of the monocyte suspension added. Plates were incubated at 37°C, 5% CO₂ until downstream analysis.

For some experiments, clusters of fibroblasts and/or monocytes were positioned in the collagen hydrogels. Cells were filtered (04-0042-2317, Sysmex), centrifuged at 300 x g for 5 min, and resuspended in 2% polyvinylpyrrolidone (PVP; P5288, Sigma-Aldrich) at 1*10^8^ or 3*10^7^ cells/mL for monocytes and fibroblasts, respectively. Nanoliter droplets of PVP/cell suspensions were injected into a 384-well micro-clear plate (781091, Greiner) pre-loaded with collagen I gels (see above), using an image-guided micro-injection robot (Life Science Methods, Leiden, NL) as described previously^40,41^. This resulted in collagen-embedded clusters with an initial diameter of 200 µm containing approximately 2000 fibroblasts or 6000 monocytes. Clusters were positioned at predefined x-y-z positions at 500 µm distance from each other. To promote monocyte migration, co-injected monocytes and fibroblasts were exposed to conditioned medium collected from fibroblasts cultured for 3 days in T75 flasks at an initial seeding density of 6 × 10⁴ cells/cm².

### Transwell assays

Monocyte chemotaxis was assessed using 3 µm pore-size transwell inserts in 24-well plates (662630, Greiner Bio-One). For migration induced by fibroblasts, 2×10⁴ fibroblasts were seeded in the lower chamber in DMEM containing 10% FBS and P/S in a final volume of 600 uL/well. Fibroblasts were incubated for 72 h with or without 10 ng/mL TGF-β1 at 37°C, 5% CO₂. Thereafter, medium was replaced with 600 µL DMEM with 2% FBS with or without 10 ng/mL TGF-β1 and incubated for another 24 h. For experiments with conditioned media, medium was collected from 3D collagen-embedded fibroblast cultures with or without 10 ng/mL TGF-β1 (see above) and added to the lower chamber in a final volume of 600 µL/well. DMEM containing 2% FBS and supplemented with 10 ng/mL CCL2 (300-04-20UG, Peprotech) was used as a positive control.

CD14+ monocytes were thawed in complete RPMI, stained with 1 µM fluorescent CellTracker Green (C2925, Invitrogen) for 20 min at 37°C, washed once with DPBS, and resuspended in DMEM containing 2% FBS at 2×10^6^ cells/mL. Where indicated, 1 µM CCR2-RA-[R] (MedChemExpress, HY-50081), an allosteric antagonist of CCR2, or 0.1% DMSO vehicle control, was added to the monocyte suspension. 100 µL of the monocyte suspension was added to the upper chamber of the transwell. Plates were incubated at 37°C, 5% CO₂, and migration was assessed by microscopy at 24 using a ZOE Fluorescent Cell Imager (Bio-Rad Laboratories). The number of migrated cells was quantified as the average number of cells per field across 10 fields in the bottom chamber of two transwells.

### RNA-seq

Fibroblasts were seeded at 6*10^4^ cells/cm^2^ and cultured for 3 days in the presence or absence of 10 ng/mL TGF-β1 in a flask. Next, cells were re-seeded at 6*10^4^ cells/cm^2^ in a flask (2D) or embedded in collagen I gel as described above and cultured for an additional 3 days in the presence or absence of 10 ng/mL TGF-β1. To collect cells for RNA isolation, 2D cultures were trypsinized and collagen-embedded cultures were treated with a blend of 1 mg/mL Collagenase and 0.1 mg/mL DNase I (B20221, Sigma-Aldrich) for 30 min at 37°C. Cells were collected in medium, counted, and cell pellets were lysed and preserved in DNA/RNA Shield buffer (R1100-50, Zymo Research). Samples were shipped to Plasmidsaurus (https://plasmidsaurus.com) for bulk RNA-seq according to the company’s workflow.

Gene-level count matrices and normalized expression values provided by Plasmidsaurus were used for downstream analysis in R (4.4.1). PCA plots and pairwise differential gene-expression analyses were performed using DESeq2 to determine the transcriptional effects of culture dimensionality and fibroblast activation. Genes with an adjusted p-value < 0.05 and an absolute log2 fold change > 1 were considered significantly differentially expressed. Genes with a log2 fold change > 1 were classified as upregulated. Genes with a log2 fold change < −1 were classified as downregulated. Gene set enrichment analysis (GSEA) was performed by Plasmidsaurus using ranked gene lists with GSEApy v0.12 using the MSigDB Hallmark gene set. Enriched pathways were evaluated based on their normalized enrichment score (NES) and false-discovery rate (FDR). Pathways with an absolute normalized enrichment score > 1.5 and an FDR-adjusted P value < 0.05 were considered significantly enriched.

### Multiplex cytokine analysis

Cytokine concentrations in conditioned medium were quantified using the LEGENDplex™ Human Essential Immune Response Panel (13-plex) with V-bottom Plate (cat. no. 740930, BioLegend), according to the manufacturer’s instructions. The panel was used to quantify IL-4, IL-2, CXCL10, IL-1β, TNF-α, CCL2, IL-17A, IL-6, IL-10, IFN-γ, IL-12p70, TGF-β1 and IL-8. Briefly, samples and standards were incubated for 2 h at RT with capture beads, followed by 1 h incubation with detection antibodies. Then, streptavidin-phycoerythrin was added and incubated for 30 min. Following washing, beads were resuspended in wash buffer and analyzed using a CytoFLEX flow cytometer (Beckman Coulter). Analyte concentrations were determined from the corresponding standard curves using LEGENDplex™ Data Analysis Software version 2025-05-01 (BioLegend).

### Immunofluorescence staining of 3D cultures

For immunofluorescence labeling of collagen I-embedded cultures, medium was removed, and hydrogels were fixed with 4% paraformaldehyde (50-00-0 VWR Chemicals) for 30 min at RT. After 3 washes with DPBS and 3 washes with DPBS containing 1% BSA (A9647, Sigma), samples were blocked and permeabilized with 1% BSA and 0.3% Triton-X-100 (9036-19-5, Sigma-Aldrich) for 1 h at 4°C on a plate shaker. Samples were stained with 1.25 µg/mL fibronectin monoclonal antibody (610077, BD Biosciences) and 1 µg/mL collagen I polyclonal antibody (PAS-95137, Invitrogen) in organoid washing buffer (OWB; DPBS containing 2% BSA, 0.1% Triton-X-100 and 0.02% SDS) for 48 h at 4 °C on a plate shaker. Samples were washed three times with OWB for 30 min and stained with secondary antibody mix containing 1 µg/mL Alexa 647 labelled goat-anti-rabbit IgG (111-605-003, Jackson), 1 µg/mL Alexa 488 labelled goat-anti-mouse IgG (A11001, Invitrogen), 1 µg/mL Hoechst 33342 (H1399, Invitrogen) and 50 nM Phalloidin Rhodamine (R415, Invitrogen) for 48 h at 4°C on a plate rocker. Finally, samples were washed three times with OWB and stored in the dark at 4°C in DPBS before imaging.

### Confocal immunofluorescence and reflection microscopy and image analysis

All confocal fluorescence images were acquired using a Nikon ECLIPSE Ti2 confocal microscope equipped with an automated stage controlled by NIS Element Software in a temperature and CO_2_ controlled incubator. Images and metadata were stored in OMERO (https://www.openmicroscopy.org/omero/).

To analyze infiltration of monocytes seeded on top of collagen gels, confocal z-stacks were acquired using Plan Fluor 10x/0.30 objective (Nikon) 48 h after monocyte seeding. Images were taken every 20 μm from the top surface of the hydrogel to 500 μm below the surface. To analyze migration of injected monocyte clusters, confocal z-stacks of images taken every 10 μm were acquired in a range of 300 μm, including the entire monocyte cluster, using a S Plan Fluor long working distance 20X/0.70 objective (Nikon) at 2 h and 24 h after injection.

Automated image analysis was performed using ImageJ (version 1.54p) image analysis software. The number and spatial coordinates of infiltrated monocytes were determined by segmenting and counting objects using the 3D Object Counter plugin^42^. For the analysis of monocyte injections, images were converted into maximum-intensity projections. A fixed-size oval region, representing the initial monocyte injection, was defined and repositioned to center the injection point for each image. CellTracker green-positive objects were identified using a fixed intensity threshold, converted into binary masks, and separated using watershed segmentation. All segmented objects located within the injection site were removed, thereby restricting the analysis to cells that had migrated beyond the initial injection area. The centroid coordinates of each remaining object were determined, and the shortest Euclidean distance from each centroid to the border of the oval region was calculated and averaged for each analyzed injection.

The collagen fiber network was visualized by confocal reflection microscopy, with a CFI Plan Apochromat LWD Lambda S 20XC water-immersion objective (Nikon) using excitation at 561 nm with a 561 nm blocking dichroic mirror for detection of the total reflection signal. Z-stack images were captured from the bottom of the well to 200 µm into the gel, with images taken at 10 μm intervals. Reflection microscopy images were shown as a maximum projection of four Z-stacks. To visualize collagen network remodeling at the upper surface of the hydrogel, confocal reflection images were captured in the top 300 µm of the hydrogel. XZ maximum-intensity projections were generated from a 10-pixel-wide region in the y-direction, corresponding to a thickness of 6.3 µm.

Automated analysis of collagen network architecture was performed using ImageJ. Because of the observed z-depth-related decrease in overall signal intensity in confocal reflection imaging, the intensity was normalized across all z-planes by normalizing the contrast, and histogram equalization was applied to each individual z-plane, allowing consistent visualization and segmentation throughout the image stack. Next, a morphological closing filter from the MorphoLibJ plugin was applied using an octagonal structuring element with a radius of 1 pixel to reduce small discontinuities and improve the continuity of detected structures^43^. Images were thresholded, pores were segmented, and the number and average size of the pores were obtained. The porosity was determined by multiplying the number and average pore size, divided by the total area.

### Statistical analyses

Statistical analysis of RNA-seq data is described in the RNA-seq section. Other statistical analyses were performed in GraphPad Prism V10.1 using paired or unpaired t-tests, or repeated-measures two-way ANOVA followed by Holm-Šídák multiple comparison test unless otherwise specified. Data were normalized for visualization, but statistical analyses were performed on unnormalized data. Statistical significance was defined as *P* < 0.05.

## Results

### TGF-β1 activation blocks fibroblast-induced monocyte infiltration of collagen I hydrogels

Chemotactic signals secreted by fibroblasts can recruit monocytes into inflamed and fibrotic tissues^14^. In addition, activated fibroblasts in such tissues remodel the ECM^6^. To address the combined effect of these properties, PBMC-derived CD14+ monocytes were incubated on top of a fibrillar collagen I hydrogel alone or loaded with skin fibroblasts (Fig 1A). Infiltration of monocytes into empty collagen gels was minimal (Fig. 1B, C). Incorporation of unstimulated fibroblasts in the collagen gels caused significant infiltration of monocytes. However, activation of the fibroblasts by exposure to TGF-β1 suppressed monocyte infiltration to values comparable to the gel-only condition.

**Figure 1:**
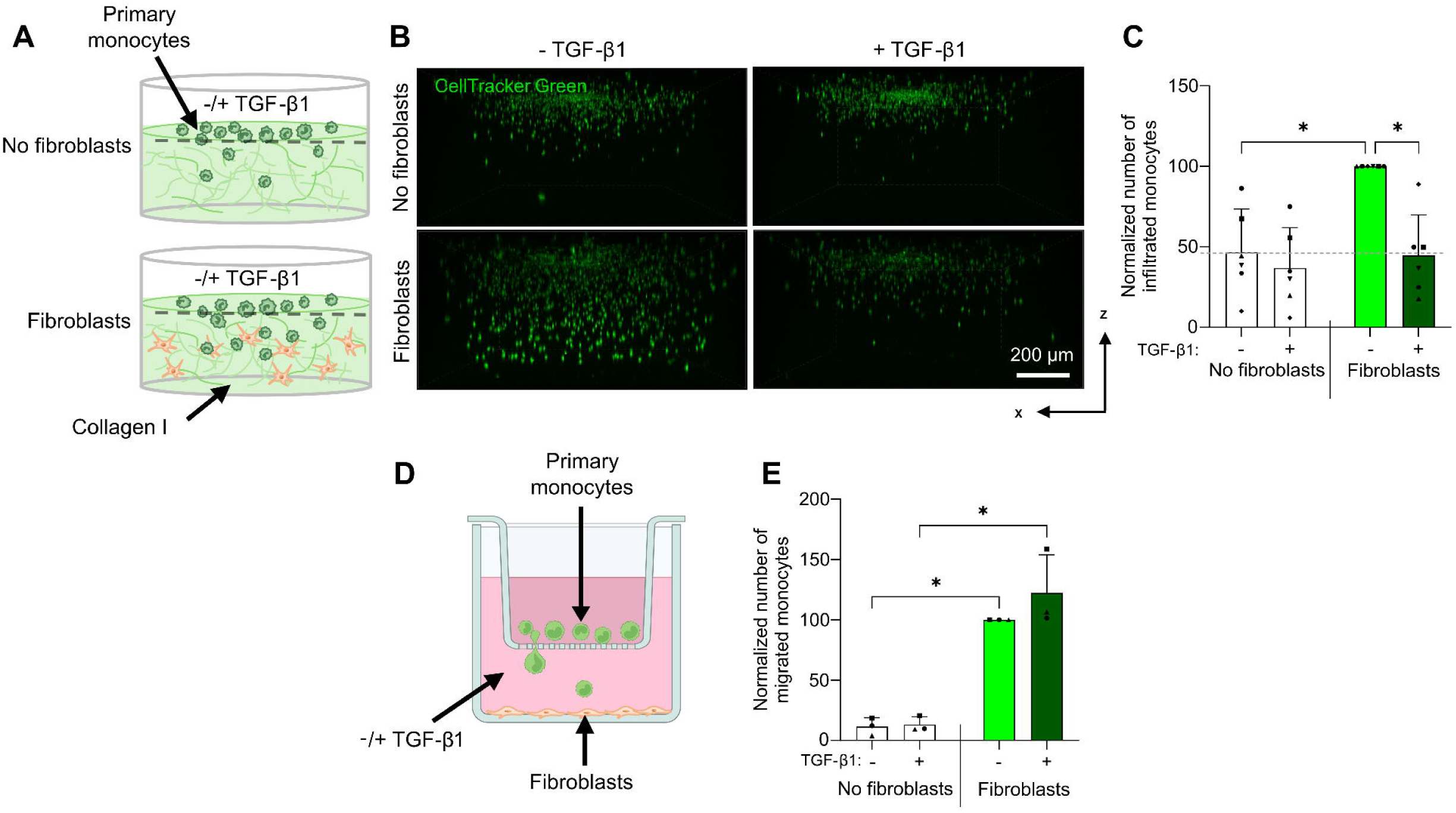
Stimulation of monocyte ECM infiltration by fibroblasts is inhibited by TGF-β1. **A**) Schematic representation of the experimental setup: fibroblasts or TGF-β1 pre-activated fibroblasts were embedded inside 2 mg/ml collagen I hydrogels in the presence or absence of 10 ng/mL TGF-β1. Fluorescent CellTracker CMFDA-labeled monocytes were seeded on top, and their infiltration into the hydrogel was assessed. Parts were generated using ChatGPT. **B**) Representative confocal fluorescent images of infiltrating monocytes in green after 48 h of co-culture. Side view max-projection. Scale bar = 200 μm. **C**) Number of infiltrated monocytes below the surface of the hydrogel after 48h, normalized to the condition using unstimulated fibroblasts. Mean ± SEM from n = 6 independent experiments (monocytes isolated from four healthy donors). Statistics: Repeated-measures two-way ANOVA followed by Holm-Šídák multiple comparison test. \**P* < 0.05. **D**) Schematic illustration of the transwell assay. Fibroblasts were seeded on the bottom of the lower chamber in the presence or absence of 10 ng/mL TGF-β1 and CellTracker CMFDA-labeled monocytes were seeded in the transwell insert. Created in https://BioRender.com **E**) Quantification of the number of monocytes that migrated towards the lower chamber, normalized to unstimulated fibroblasts. Mean ± SEM from n = 3 independent experiments (monocytes isolated from three healthy donors). Statistics: repeated-measures two-way ANOVA followed by Holm-Šídák corrected multiple comparisons test. \**P* < 0.05.

To assess if the recruitment was mediated by chemotactic signals secreted by fibroblasts and whether this was affected by TGF-β1, monocyte recruitment was studied in a transwell assay (Fig. 1D). Migration of monocytes towards medium only was negligible, while a 2D culture of fibroblasts grown in the bottom chamber led to significant migration of monocytes (Fig.1E). In this setup, stimulation of fibroblasts with TGF-β1 did not reduce monocyte attraction. Notably, TGF-β1 did not affect monocyte migration in absence of fibroblasts in the transwell or monocyte infiltration of empty collagen I gels, excluding a direct effect on monocytes.

These results showed that TGF-β1 activation attenuated the ability of fibroblasts to stimulate monocyte infiltration into collagen hydrogels, whereas TGF-β1 did not affect chemotactic monocyte attraction by fibroblasts per se.

### TGF-β1 activation induces a switch from inflammatory signaling to ECM remodeling signatures in collagen I-embedded fibroblasts

To investigate how culture dimensionality and TGF-β1 exposure affected fibroblasts, RNA-seq was performed (Fig. 2A). Both culture dimensionality and TGF-β1 stimulation induced substantial changes in gene expression, resulting in large numbers of differentially expressed genes (DEGs) (Fig. S1A). 1000 genes were up- and 537 genes were downregulated in unstimulated 3D collagen-embedded versus 2D cultures. TGF-β1 stimulation in 2D resulted in 520 up- and 967 downregulated genes. In 3D collagen-embedded cultures, 522 genes were up- and 1164 genes were downregulated in response to TGF-β1. Principal component analysis (PCA) showed that samples from the same experimental condition clustered together, with separation according to TGF-β1 treatment along PC1 (Fig. 2B). The genes with the highest positive loadings on PC1 included markers related to inflammation (*PTX3, ACKR4, SECTM1, CCL2, IL1R1,* and *CEMIP*) and ECM interaction (*COL12A1, DCN, MMP11*) (Fig. S1B). Negative-loading genes were related to ECM remodeling (*POSTN, HAPLN1, ELN, COL4A1, CCN2,* and *TGFBI)*. Along PC2, 2D and 3D culture samples were separated, with genes associated with ECM organization and fibroblast activation (*COL10A1, MMP11, POSTN, MFAP4, LTBP2,* and *TGFBI*) contributing to the variance in PC2 (Fig. S1C).

**Figure 2.**
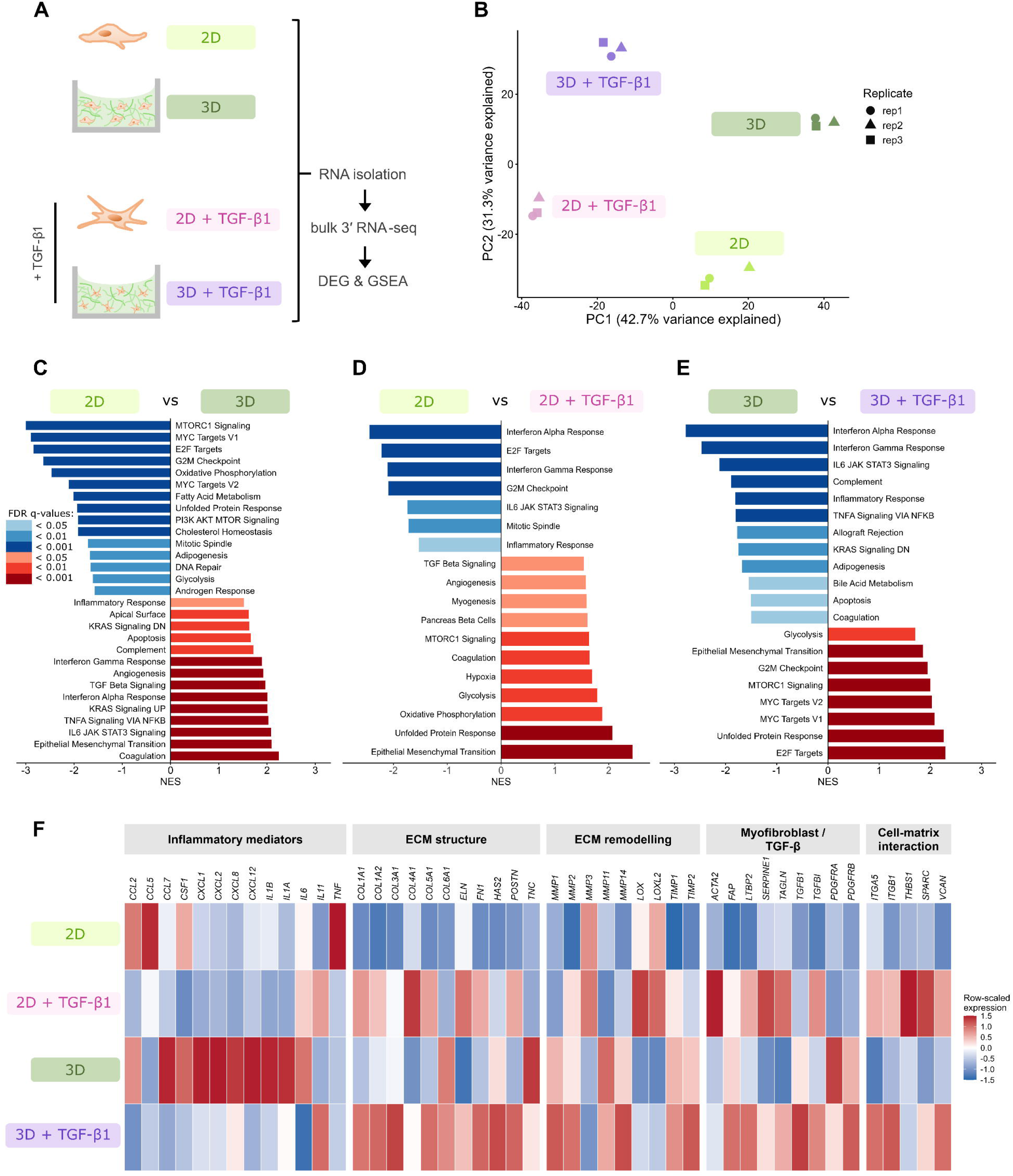
RNA-seq analysis reveals transcriptional differences in inflammatory profile and ECM remodeling induced by 3D collagen culture and TGF-β1 exposure. **A**) Schematic overview of the experimental conditions and RNA sequencing workflow. Fibroblasts or pre-activated fibroblasts were cultured in 2D or embedded in 3D in 2 mg/mL collagen I hydrogels, in the presence or absence of 10 ng/mL TGF-β1 for 3 days, followed by RNA isolation, DEG analysis, and GSEA. **B)** PCA of the four conditions with three biological replicates, showing clustering of the replicates and separation of the experimental conditions. **C-E)** GSEA-derived pathways ranked by NES. Pathways are shown if |NES| > 1.5 and FDR q-value < 0.05. Colors indicate FDR significance categories as defined in the plot legend. **C)** Enriched gene sets in 2D (blue) versus enriched in 3D (red). **D)** Enriched gene sets in 2D - TGF-β1 (blue) versus enriched in 2D + TGF-β1 (red). **E)** Enriched gene sets in 3D - TGF-β1 (blue) versus enriched gene sets in 3D + TGF-β1 (red). **F)** Heatmap of row-scaled expression for manually selected genes, grouped across the fibroblast culture conditions. Raw counts were variance-stabilized using DESeq2, averaged across three biological replicates per condition, and row-scaled per gene. Only genes detected after low-expression filtering were included. Color indicates relative expression across conditions for each gene, with blue representing lower and red representing higher relative expression.

Unbiased GSEA of the RNA-seq data showed that resting fibroblasts cultured in 2D were enriched for cell cycle, metabolic, and biosynthetic programs, whereas 3D collagen-embedded resting fibroblasts were enriched for inflammatory, cytokine-response, and epithelial-mesenchymal-transition (EMT) programs (Fig. 2C). In 2D fibroblast cultures, exposure to TGF-β1 enriched metabolic, developmental and EMT programs accompanied by a decrease in inflammatory and cell cycle related enrichment (Fig. 2D). Exposure to TGF-β1 in 3D cultures suppressed the inflammatory and cytokine-response signatures and instead caused a coordinated metabolic rewiring to a glycolytic state with enrichment of cell-cycle and EMT programs (Fig. 2E).

We next selected a set of genes (Table S1) to evaluate patterns in transcriptional changes related to inflammation, ECM structure, ECM remodeling, myofibroblast differentiation, and cell-matrix interaction in 2D and 3D collagen-embedded fibroblast cultures. Row-scaled expression in the heatmaps showed distinct, condition-dependent expression patterns among the gene sets (Fig. 2F). Notably, some candidate genes, such as *IFNG*, *IL-10*, and *MMP9*, were not retained in the heatmap due to low or undetectable counts in the RNA-seq dataset. In 2D cultures, TGF-β1 had little effect on the expression of inflammatory mediators (except for a significant downregulation of *CCL2 and CSF1*). By contrast, the highly expressed inflammatory mediators in 3D were strongly downregulated in response to TGF-β1. A moderate upregulation of genes associated with ECM structure and ECM remodeling was observed in both 2D and 3D cultures exposed to TGF-β1. Genes associated with myofibroblast phenotype/TGF-β1 response and cell-matrix interaction were moderately upregulated in both 2D and 3D collagen I-embedded fibroblast cultures upon TGF-β1 stimulation.

These data demonstrated that shifting from 2D culture to 3D collagen-embedded fibroblast culture was accompanied by a shift toward a more cytokine-responsive, inflammatory state. This shift also led to a distinct response to TGF-β1: in 2D, TGF-β1 increased ECM interaction, ECM remodeling, and myofibroblast phenotype, whereas in 3D collagen-embedded fibroblast cultures, it strongly suppressed inflammatory mediators and more robustly enhanced ECM remodeling capacity.

### TGF-β1 exposure suppresses the inflammatory secretome of collagen-embedded fibroblasts, which partially inhibits monocyte attraction

To further explore the possibility that downregulation of inflammatory chemokines/cytokines could explain the fact that TGF-β1 suppressed the ability of fibroblasts to trigger monocyte infiltration in 3D collagen matrices, we analyzed secretion of a panel of cytokines and chemokines. Out of 13 soluble analytes, IFN*γ*, IL-10, IL-12p70, IL1β, IL-2, IL-4, CXCL10, IL- 17A, and TNF*a* were not detected (Fig. S2). IL-6 and IL-8 were detected at low levels in medium from 2D cultured fibroblasts and moderately affected by TGF-β1, while CCL2 levels were higher but completely suppressed by exposure to TGF-β1 (Fig. 3A; Fig. S2). In media from collagen-embedded fibroblast cultures, IL-6, IL-8, and CCL2 were detected and, in agreement with the transcriptional analysis (Fig. 2F), TGF-β1 treatment strongly suppressed the levels of all three chemokines. Lastly, free active TGF-β1 was readily detected in TGF-β1-exposed 2D cultures but absent from media derived from TGF-β1-exposed collagen-embedded cultures, indicating its trapping in the collagen scaffold (Fig. S2).

**Figure 3:**
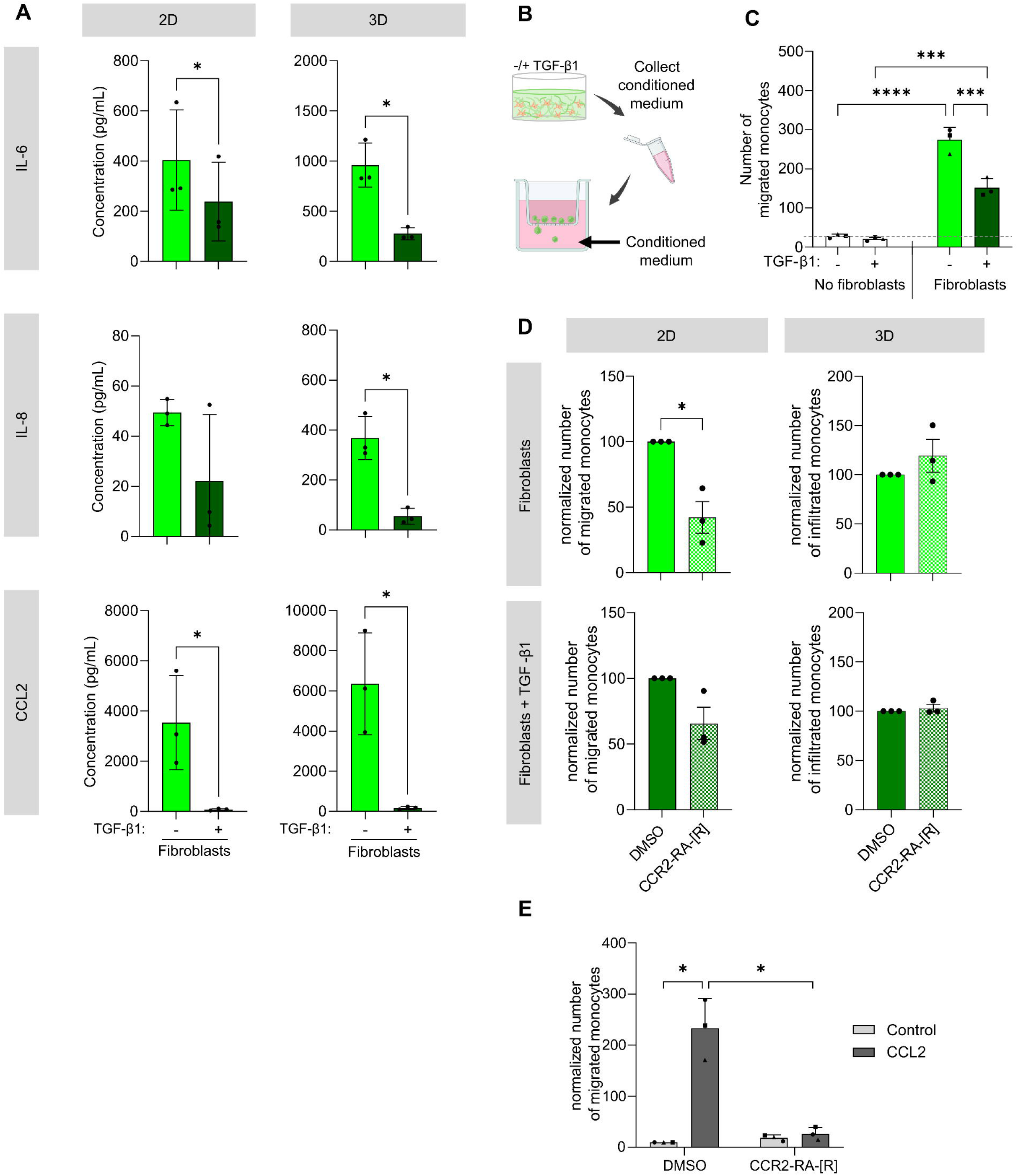
Fibroblast activation decreases the secretion of IL-6, IL-8 and CCL2 and partially inhibits chemotactic attraction of monocytes. **A)** The concentration of IL-6, IL-8 and CCL2 in medium collected from fibroblasts cultured in 2D or 3D was measured by bead-based multiplex immunoassay. Mean ± SD from n = 3 independent experiments. Statistics: unpaired t-test. *P < 0.05. **B)** Schematic overview of conditioned medium collection and subsequent transwell assay. Created in https://BioRender.com **C)** Quantification of the number of monocytes that migrated towards the lower chamber filled with conditioned medium from 3D collagen I-embedded fibroblasts cultured in the presence or absence of TGF-β1. Mean ± SD for medium collected from n = 3 independent experiments (monocytes isolated from one healthy donor). Statistics: Two-way ANOVA, followed by Holm-Šídák corrected multiple comparison test. \*\*\**P* < 0.001; \*\*\*\**P* < 0.0001. **D**) The effect of 1 µM CCR2-RA-[R] on monocyte attraction by 2D cultures of fibroblasts or TGF-β1-activated fibroblasts in transwell setup, and infiltration of monocytes into 3D collagen I-embedded fibroblasts or TGF-β1-activated fibroblasts. Data is normalized to the DMSO-treated vehicle condition. Mean ± SEM, with n = 3 independent experiments (monocytes isolated from two healthy donors). Statistics: Paired t-test \**P* < 0.05. **E)** Quantification of the number of monocytes that migrated towards medium containing 10 ng/ml CCL2 in the lower chamber of a transwell setup in the absence or presence of 1 µM CCR2-RA-[R], normalized to DMSO control. Mean ± SEM with n = 3 independent experiments (monocytes isolated from two healthy donors). Statistics: Repeated measures two-way ANOVA with Holm-Šídák corrected multiple comparisons. \**P* < 0.05.

We next asked whether conditioned medium derived from collagen-embedded fibroblast cultures attracted monocytes and how this was affected by TGF-β1 (Fig. 3B). Monocytes migrated toward conditioned medium from collagen-embedded fibroblasts, confirming that these fibroblasts secrete soluble monocyte-attracting factors. TGF-β1 activation of the collagen-embedded fibroblasts reduced the monocyte attraction capacity of conditioned medium by ∼50% compared to unstimulated fibroblasts (Fig. 3C).

CCL2 can mediate monocyte recruitment by fibroblasts in inflamed tissue^18^. The fact that CCL2 secretion by collagen-embedded fibroblast cultures was abrogated in response to TGF-β1 exposure (Fig. 3A) while the conditioned medium retained ∼50% of its monocyte-attracting potential (Fig. 3C), suggested CCL2 was not, or not solely, involved. To test this hypothesis, the effect of CCR2-RA-[R], a degrader of the CCR2 receptor for CCL2, was evaluated. CCR2-RA-[R] partially inhibited monocyte recruitment by 2D cultured fibroblasts in the transwell assay, while it did not affect fibroblast-induced monocyte infiltration into collagen gels (Fig. 3D). As a positive control, the same concentration of CCR2-RA-[R] completely blocked monocyte migration that was induced by recombinant CCL2 (Fig. 3E).

These results showed that reduced secretion of inflammatory cytokines and chemokines can partially explain the suppression of fibroblast-induced monocyte infiltration into collagen gels by TGF-β1 but other aspects must be involved.

### Inhomogeneous ECM remodeling by TGF-β1-activated fibroblasts

To further explore the emergence of an ECM remodeling phenotype in collagen-embedded fibroblast cultures upon TGF-β1-exposure (Fig. 2F), we investigated how TGF-β1 treatment of fibroblasts structurally affected the ECM fiber network (Fig. 4A). Exposure of fibroblasts to TGF-β1 led to an increased F-actin cytoskeleton with more prominent actin filament bundles, and densification of the nearby ECM (Fig. 4B, C). Moreover, confocal reflection microscopy showed that inclusion of fibroblasts increased porosity of the ECM fiber network, and larger pores formed in TGF-β1-exposed collagen-embedded fibroblast cultures (Fig 4B, D). This suggested strong contractile activity of TGF-β1-stimulated fibroblasts leading to rearrangement of the fiber network. Indeed, inhibition of contractility by treatment with the myosin II inhibitor blebbistatin suppressed ECM remodeling (Fig. S3). In addition to contractile remodeling, staining for human collagen I and fibronectin showed collagen I production and deposition of fibronectin fibrils upon TGF-β1 treatment (Fig. 4E). Notably, TGF-β1 led to strong densification of the ECM at the gel-liquid interface of the upper region of the fibroblast-containing collagen hydrogel (Fig. 4F).

**Figure 4:**
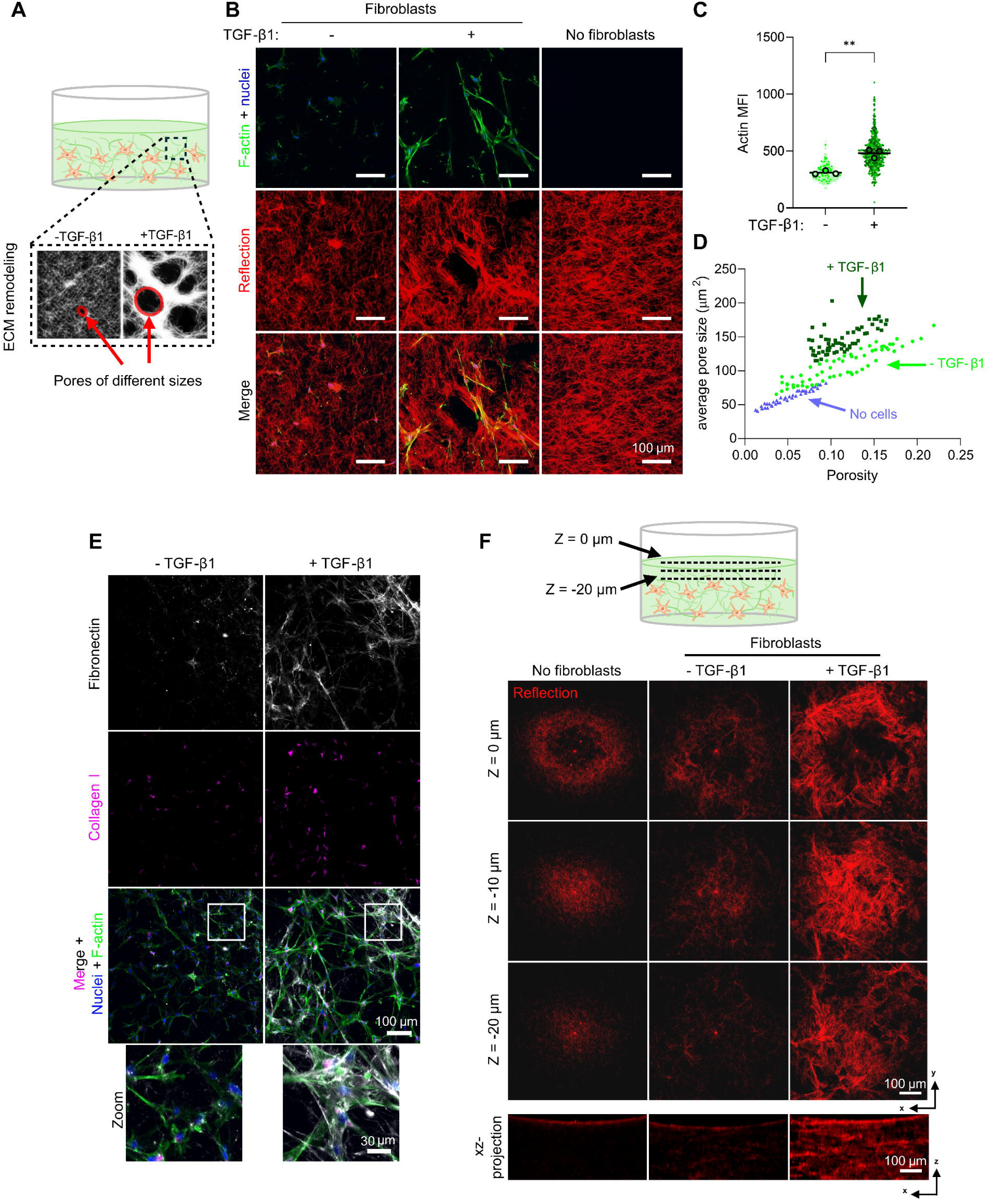
Fibroblasts increase 3D collagen network porosity and TGF-β1 activation causes ECM densification at the hydrogel liquid interface. **A**) Schematic illustration of analysis of ECM remodeling by fibroblasts. Red circles indicate pores within the ECM network. **B)** Representative maximum-intensity projections of confocal microscopy imaging showing collagen I gels with or without fibroblasts or TGF-β1-activated fibroblasts. F-actin (Phalloidin iFluor 488, green), nuclei (Hoechst, blue), ECM fibers detected by reflection microscopy (red). Scale bar = 100 μm. **C**) Quantification of mean fluorescence intensity (MFI) of the F-actin cytoskeleton. Dots represent single-cell measurements. Welch’s *t*-test was performed on the average MFI of all cells measured in three independent experiments, indicated by black circles. \*\**P* < 0.01. **D**) Scatter plot showing average pore size and overall porosity quantified from reflection microscopy images as shown in B. Dots represent single-cell measurements obtained from three independent experiments. **E**) Maximum-intensity projections of confocal microscopy images showing immunofluorescence staining of fibronectin (white), collagen I (pink), or an overlay of both with F-actin (Phalloidin iFluor 488, green) and nuclei (Hoechst, blue) for collagen gels containing fibroblasts cultured in the absence or presence of TGF-β1. Scale bar = 100 μm. White boxes indicate the zoomed-in area shown at the bottom. Scale bar = 30 μm. **F**) Reflection microscopy images acquired top z-planes (0 μm, -10 μm, -20 μm) and xy-maximum-intensity projections illustrating densification of collagen fiber network at the hydrogel liquid interface by TGF-β1-stimulated fibroblasts. Scale bar = 100 μm.

These results showed that TGF-β1-stimulated fibroblasts strongly remodeled their surrounding microenvironment through contraction and ECM protein secretion. This resulted in large pores and densified areas within the ECM, including marked densification at the hydrogel-liquid interface.

### TGF-β1 activation does not interfere with overall fibroblast-induced monocyte migration within the collagen I network

Since fibroblast-induced monocyte infiltration of fibrillar collagen hydrogels was abrogated in the presence of TGF-β1 (Fig. 1C) while chemotactic attraction of monocytes under those conditions was only partially inhibited (Fig. 3C), we considered ECM remodeling as an additional underlying mechanism. First, we tested whether the densified collagen I fiber network caused by TGF-β1-stimulated fibroblasts at the surface of the hydrogels (Fig. 4F), rather than global ECM remodeling, represented a local barrier. For this purpose, small clusters of monocytes were injected into empty- or fibroblast-containing hydrogels to bypass the potential barrier caused by TGF-β1-stimulated fibroblasts at the hydrogel-liquid interface (Fig. 5A). Migration of monocytes in empty collagen gels was minimal (Fig. 5B). Inclusion of fibroblasts stimulated migration, as analyzed by calculating the number of migrating monocytes and their average covered distance (Fig. 5C,D). Interestingly, in this experimental setup (in contrast to results with monocytes seeded on top of the collagen I hydrogels (Fig. 1C)), fibroblast-induced monocyte migration was not affected by TGF-β1 activation. The use of fibroblast-conditioned medium also stimulated monocyte migration, albeit to a lesser extent and, again, this effect was not suppressed by TGF-β1 activation.

**Figure 5:**
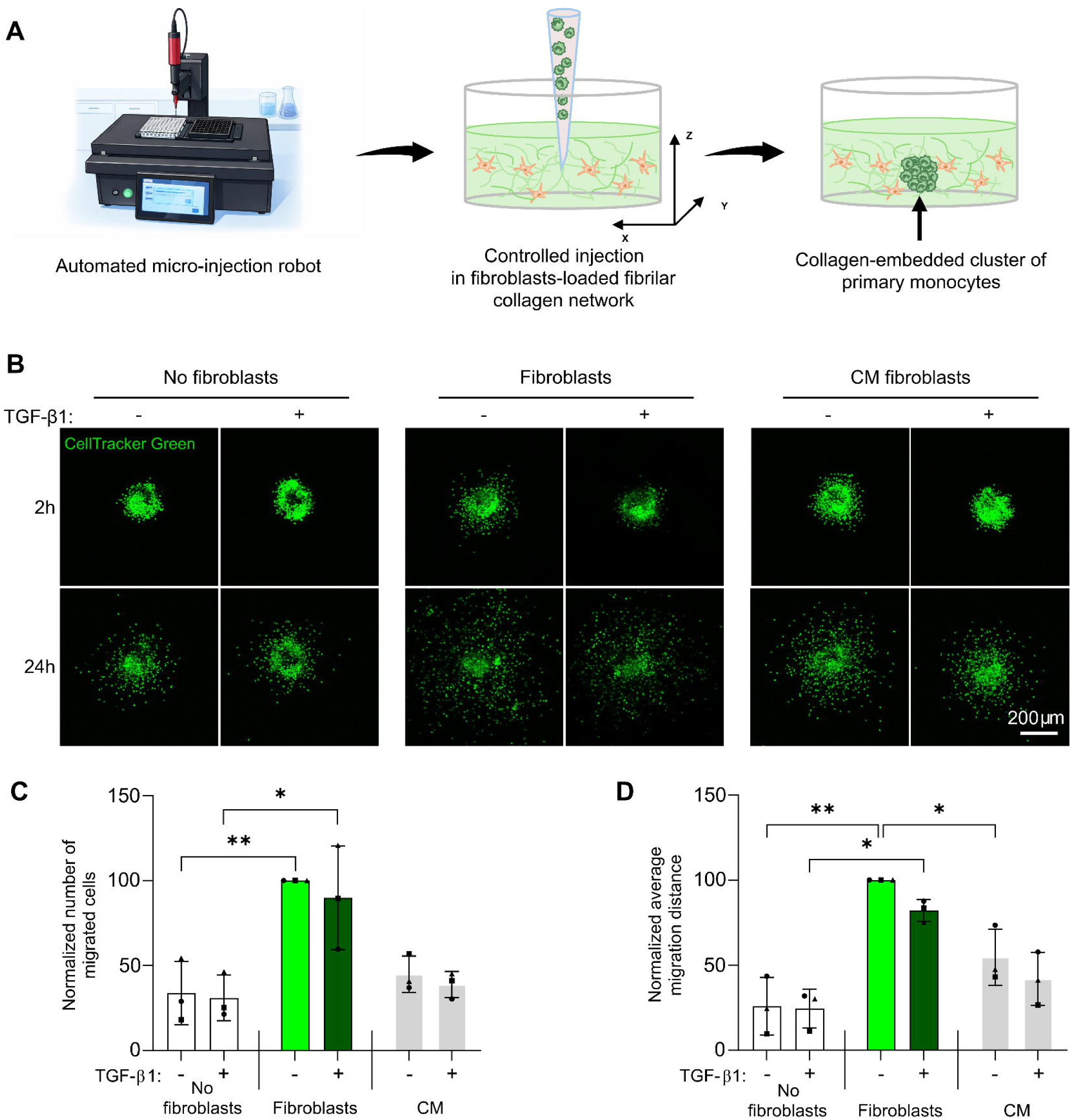
Monocyte migration induced by fibroblasts is not affected by TGF-β1 when monocytes are injected into the gel. **A)** Schematic representation of the experimental setup, in which defined clusters of fluorescent CellTracker Green-labeled monocytes are injected inside collagen I gels containing fibroblasts in the presence or absence of TGF-β1. **B)** Representative maximum-intensity projections of confocal images of monocytes (green) inside empty hydrogels in the presence or absence of TGF-β1, hydrogels containing fibroblasts in the presence or absence of TGF-β1, or hydrogels treated with conditioned medium (CM) harvested from fibroblasts or TGF-β1 stimulated fibroblasts. Images were acquired at 2 h and 24 h after injection of the monocyte clusters. Fibroblasts are not labeled. Scale bar = 200 μm. **C)** Quantification of the number of monocytes that had migrated at t = 24h outside the cluster border at t = 2h. Data is normalized to the untreated fibroblast condition. **D)** Quantification of the average migration distance for all monocytes analyzed in C. Bars show the mean ± SD of three independent experiments with monocytes isolated from two different healthy donors. Statistical analysis for C and D was performed using repeated measures two-way ANOVA followed by Holm-Šídák multiple comparison test. *P < 0.05; **P < 0.01.

These results indicated that fibroblast-induced migration of monocytes within a collagen I network is not affected by TGF-β1 but, instead, the changes at the hydrogel-liquid interface caused by TGF-β1-stimulated fibroblasts prevent monocyte entry.

### Excessive local ECM remodeling by clusters of TGF-β1-stimulated fibroblasts restricts monocyte migration

Fibroblasts *in vivo* have been shown to cluster and show collective behavior, particularly in pathologies such as fibrosis or cancer^44,45^. We previously reported how clusters of fibroblasts can cause extensive ECM remodeling^41^. To test whether such remodeling can also lead to local inhibition of monocyte migration, various patterns of TGF-β1-stimulated fibroblast clusters were designed in the collagen hydrogels by microinjection, and the fate of monocytes in their vicinity was monitored. A single cluster of fibroblasts caused no visible ECM remodeling two days after injection (Fig. 6A), and monocytes subsequently placed in its vicinity (at a distance of 250 µm) preferentially moved in the direction of the fibroblast cluster (Fig. 6B-C). Two clusters of fibroblasts spaced 500 µm apart created a bridge of densified ECM after two days (Fig. 6A). For monocytes subsequently placed in the middle of this densified area between the fibroblast clusters, migration was attenuated (Fig. 6D, E,). Lastly, four clusters of fibroblasts spaced 500 µm apart created strongly densified ECM bridges between the corners, forming a square (Fig. 6A). Monocytes subsequently placed in the center of the square were trapped as compared to monocytes not surrounded by fibroblast clusters, showing almost no monocytes passing through the strongly remodeled ECM (Fig. 6F,G).

**Figure 6:**
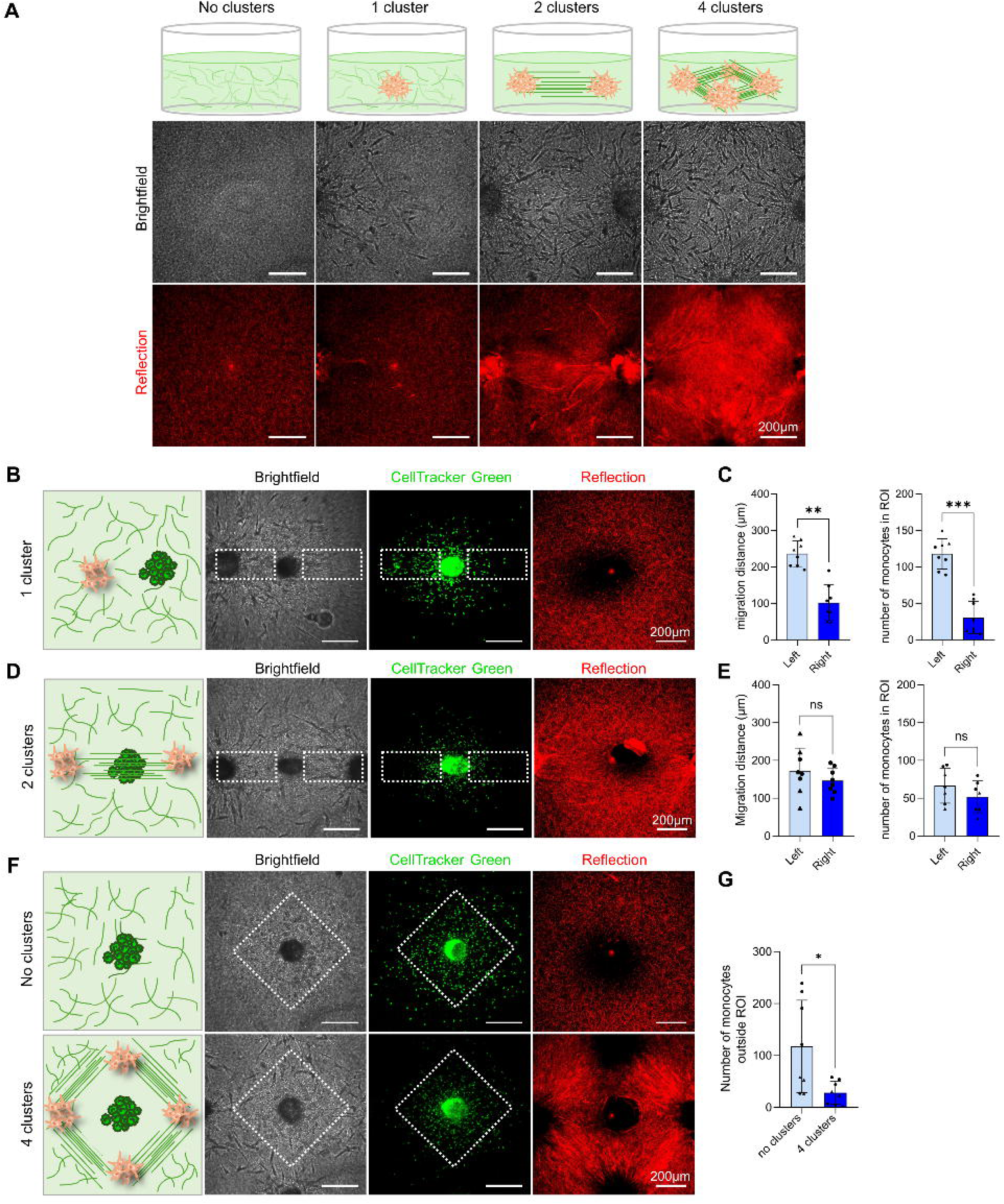
Local ECM remodeling by fibroblasts blocks monocyte migration. **A)** Schematic cartoon of patterns of injected clusters of TGF-β1-activated fibroblasts and representative brightfield and maximum intensity projection confocal reflection microscopy images. Scale bar = 200 µm. **B,D)** Co-injections of clusters of TGF-β1-activated fibroblasts (unlabeled) and clusters of monocytes (CellTracker green-labeled). Reflection microscopy in red showing ECM remodeling by one or two fibroblast clusters. The white boxes indicate the regions of interest (ROIs) used for analysis. Scale bar = 200 µm. **C**) Quantification of the number of monocytes and the average migration distance from the injection core in the left and right ROI when monocytes are co-injected with one fibroblast cluster. **E**) Quantification of the number of monocytes and their average migration distance from the injection core in the left and right ROI when monocytes are co-injected with two fibroblast clusters. In C and E, bar graphs show the mean ± SD from two independent experiments, indicated by dots and triangles, with a total of 7-8 replicates. Comparisons between matched ROIs were performed using a paired t-test. ns, non-significant; \*\**P* < 0.01; \*\*\**P* < 0.001. **F**) Schematic cartoon and representative brightfield and maximum intensity projection confocal microscopy images of one cluster of monocytes (CellTracker green-labeled) alone or surrounded by four injected clusters of TGF-β1-activated fibroblasts (unlabeled). Reflection microscopy in red shows that four fibroblast clusters strongly remodel the ECM. The white boxes indicate the ROIs used for analysis. **G**) Quantification of the number of monocytes outside the ROI in the absence or presence of four fibroblast clusters. The bar graph represents the mean ± SD from two independent experiments with a total of 7-8 replicates. Statistical analysis was performed by an unpaired t-test. \**P* < 0.05.

These data demonstrated that local ECM remodeling by TGF-β1-stimulated fibroblast clusters can create barriers that locally restrict monocyte migration.

## Discussion

The interplay between fibroblast-driven chemotactic signaling and ECM remodeling, and how these processes jointly regulate monocyte infiltration in fibrotic tissues, remains unresolved. Our combined results show that fibroblasts embedded in a 3D fibrillar collagen network promote i) monocyte entry into and ii) migration through an ECM network by adopting an inflammatory phenotype and producing chemotactic signals. In addition, fibroblasts increase the porosity of the collagen fiber network. TGF-β1 activation of the fibroblasts suppresses inflammatory transcriptional programs and secretion of inflammatory mediators. Instead, the activated fibroblasts switch to an ECM-producing and ECM-remodeling phenotype. The resulting reduction in chemotactic signals and enhanced matrix remodeling does not affect overall fibroblast-induced monocyte migration within the collagen fiber network. However, ECM densification at hydrogel-liquid interfaces or near clusters of TGF-β1-activated fibroblasts prevents tissue infiltration and locally restricts monocyte migration. Together, these findings support a model in which inflammatory fibroblasts stimulate monocyte tissue infiltration, but their strong activation in fibrotic tissues decreases recruitment and creates barriers leading to exclusion of monocytes.

The use of advanced 3D co-culture models together with confocal microscopy techniques, RNA-seq and secretome analysis allowed us to study monocyte migration, fibroblast activation, spatial organization, and ECM remodeling. These dynamic processes are particularly difficult to study *in vivo* and highlight the value of advanced 3D *in vitro* models for investigating immune-stromal interactions in fibrosis^46^.

Our finding that transitioning dermal fibroblasts from 2D to 3D activates inflammatory signaling is in line with a previous study showing that pancreatic CAFs cultured in 2D exhibit a more myofibroblast phenotype, while 3D CAF spheroids obtain an inflammatory state that promotes chemotactic recruitment of monocytes^47^. Moreover, others have reported enrichment of ECM-remodeling pathways in a 3D dermal fibroblast sheet model or lung fibroblasts in 3D collagen as compared to the corresponding 2D cultures^48,49^. In agreement, we find induction of myofibroblast and ECM-associated genes such as *FAP*, *TGFB1*, *MMP14*, *HAS2*, and *TNC* in 3D culture but in our study the induction of inflammatory mediators such as *CXCL2, CXCL8 CXCL12* and *IL1B* is particularly prominent. Indeed, conditioned medium from collagen-embedded fibroblasts chemotactically recruits monocytes, which is in line with other reports that connect fibroblasts to monocyte infiltration^27,50–52^. These studies implicate CCL2 in this process, and using CCR2-RA-[R] we indeed show that CCL2 signaling is involved in fibroblast-mediated monocyte attraction. However, infiltration of collagen-embedded fibroblast cultures is not affected by CCR2-RA-[R], indicating that other mediators, including the detected IL-8 and/or chemokine signaling through CCR1 and CCR5, may contribute to monocyte recruitment^53^. CCL2-dependent chemotaxis is likely embedded within a broader network of biochemical and biophysical cues, and its impact context-dependent.

The observation that fibroblasts stimulate monocyte migration through ECM aligns with earlier reports. Tissue infiltration by monocytes involves their navigation through the collagen I-rich ECM, and fibroblasts can substantially alter the ECM architecture through which migration occurs^28,54^. ECM stiffening by activated fibroblasts^47^ has been shown to promote monocyte migration^28^. Also, fiber alignment, tunnel formation, and dynamic contractility caused by fibroblasts promote monocyte migration in a collagen meshwork^29,30^. We find that scattered fibroblasts in 3D collagen I cause increased meshwork porosity, which can contribute to monocyte migration. Moreover, fibroblast-induced monocyte migration within the 3D collagen network can be explained by ECM binding of pro-migratory factors^55^, direct cell-cell contact with fibroblasts^56^, or favorable biomechanical alterations in the microenvironment^28^. The enhanced infiltration and migration of monocytes in fibroblast-loaded collagen gels will likely be a combined effect of soluble signaling and biomechanical matrix properties.

Our RNA-seq data show a striking switch from inflammatory signaling to ECM remodeling upon TGF-β1 activation of 3D collagen-embedded fibroblasts. Analyses of the secretome and imaging of the ECM architecture support this switch. The finding that the response to TGF-β1 differs for 2D versus 3D collagen-embedded fibroblast cultures extends a previous observation that culture dimensionality shapes the response of MRC5 lung fibroblasts to TGF-β1^57^.

Fibroblasts are increasingly recognized as a heterogeneous and highly plastic cell population that can adopt distinct functional states depending on tissue context and environmental cues. Single-cell transcriptomic studies have identified fibroblast populations specialized in immune-regulatory and inflammatory programs, alongside states predominantly associated with ECM production, contraction, and tissue remodeling^15,58^. The inflammatory phenotype induced by 3D culture in our model and the switch to ECM remodeling induced by TGF-β1 support the concept that fibroblast function is strongly context-dependent and that inflammatory and matrix-remodeling activities can represent distinct fibroblast states. Moreover, temporal human wound data have revealed that the functional state of fibroblasts changes during wound healing. Inflammatory myofibroblasts dominated early, whereas the predominant population in late-stage wounds had shifted to a myofibroblast state^59^. The pronounced shift from an inflammatory to an ECM-producing and ECM-remodeling phenotype we identify supports the concept that inflammatory and matrix-remodeling fibroblast programs are separate and dynamically regulated.

The enhanced ECM remodeling capacity of TGF-β1-activated fibroblasts leads to the appearance of large holes and areas with local densification of the collagen meshwork. This can be attributed to contractility-driven inhomogeneous remodeling, and we find it does not affect global monocyte migration within the fiber network. Monocytes are highly motile cells that use amoeboid migration to crawl through heterogeneous extracellular spaces^60^. The changes caused by scattered activated fibroblasts likely change the paths of monocytes as they navigate through regions of lower physical resistance without inhibiting migration, as has been described for leukocytes^61–63^. Thus, ECM remodeling by individual, spatially dispersed fibroblasts may alter local migration routes without imposing a global barrier to monocyte movement through the matrix.

However, when physical tissue constraints exceed the capacity of a cell to deform, migration is limited^64^. We find that extensive collagen network remodeling caused locally when activated fibroblasts form clusters creates such barriers, obstructing monocyte migration. This reflects collective ECM remodeling that has also been observed *in vivo* for fibroblasts at sites of tissue damage^38^, cardiac infarction^39^, and IPF^44^. Our findings indicate such regions can locally restrict monocyte infiltration. Moreover, we observe that local densification at the ECM-liquid interface appearing at the surface of hydrogels containing scattered activated fibroblasts also forms a barrier that prevents entry of monocytes into the ECM network. A similar mechanism has been recently hypothesized by others^65^. Together, these observations provide a potential mechanistic explanation for spatial distribution and immune-restricted fibrotic niches observed in human disease. In IPF, fibroblastic myofibroblast-rich lesions can be found surrounded but not penetrated by immune-cell infiltrates, and fibroblastic foci with low CD45+ immune-cell abundance have been described as “cold” lesions with a more progressive fibrotic transcriptional profile^31,37^. Similarly, in renal fibrosis, myofibroblast-rich glomerular regions can display low densities of T-cells, B-cells, and macrophages despite a strong profibrotic signature^32^. Comparable spatial restriction is well established in desmoplastic tumors, where fibroblast-rich regions and dense or aligned collagen networks can restrict immune cell access and migration^35,66^. Thus, while fibroblast activation does not globally suppress monocyte migration within the ECM, the spatial heterogeneity of fibroblast density and ECM remodeling can generate local physical barriers that restrict monocyte access to specific fibrotic regions.

Altogether, our work sheds new light on how the intricate balance between paracrine signaling and architectural organization of the ECM by fibroblasts shapes monocyte infiltration. Fibroblasts scattered through the ECM produce chemotactic factors and enhance the porosity (permissiveness) of the ECM, thereby promoting monocyte infiltration. This is consistent with an inflammatory and tissue-remodeling environment where fibroblasts can facilitate immune-cell recruitment and migration. TGF-β1 activation, however, shifts fibroblasts toward an ECM- remodeling phenotype while reducing the expression of chemotactic factors, which may attenuate monocyte recruitment. Moreover, enhanced ECM remodeling generates barriers at ECM–liquid interfaces that prevent monocyte entry and, when fibroblasts accumulate locally, produces highly remodeled regions that restrict monocyte migration. Our findings therefore suggest that increasing fibroblast activation and local fibroblast accumulation can shift the microenvironment from one that promotes monocyte recruitment and infiltration towards one in which reduced chemotactic signaling and spatially restricted ECM barriers limit immune-cell accessibility. This provides a mechanistic basis for the reported spatial distribution of monocytes and emergence of immune-poor or “cold” fibrotic niches observed in human fibrotic tissues.

## Acknowledgements

The authors thank Dr. Natalia Ortiz Zacarias, Prof. dr. Laura H. Heitman for providing the CCR2-RA-[R] antagonist, Ms. Jasmijn H.M. Hundscheid for assisting with experimental work, and the Leiden Cell Observatory for assistance in this work.

This work was supported by the SMART Organ-on-Chip Perspectief Program of the Dutch Research Council (NWO, project number P19-03).

## Data availability

Raw FASTQ files and processed data are deposited in BioStudies. and are accessible through the number S-BSST3507. The RNA-seq analysis pipeline used in this paper is documented at https://doi.org/10.5281/zenodo.23670017

Any additional information regarding the data reported in this paper is available from the contact person upon request.

## Author contributions

SdW performed experiments, analyzed data, and drafted the manuscript. EN supervised the study and critically revised the manuscript. EHJD conceived and supervised the study, acquired funding, and critically revised the manuscript. All authors read and approved the final manuscript.

## Declaration of generative AI and AI-assisted technologies in the writing process

During the preparation of this work, the author(s) used ChatGPT (OpenAI) in order to assist with language editing and improving textual structure of certain sections of the manuscript, assist with coding in R, and for generation of a schematic conceptual illustration. After using this tool/service, the author(s) reviewed and edited the content as needed and take(s) full responsibility for the content of the published article.

---

**Supplementary Figure 1 :**
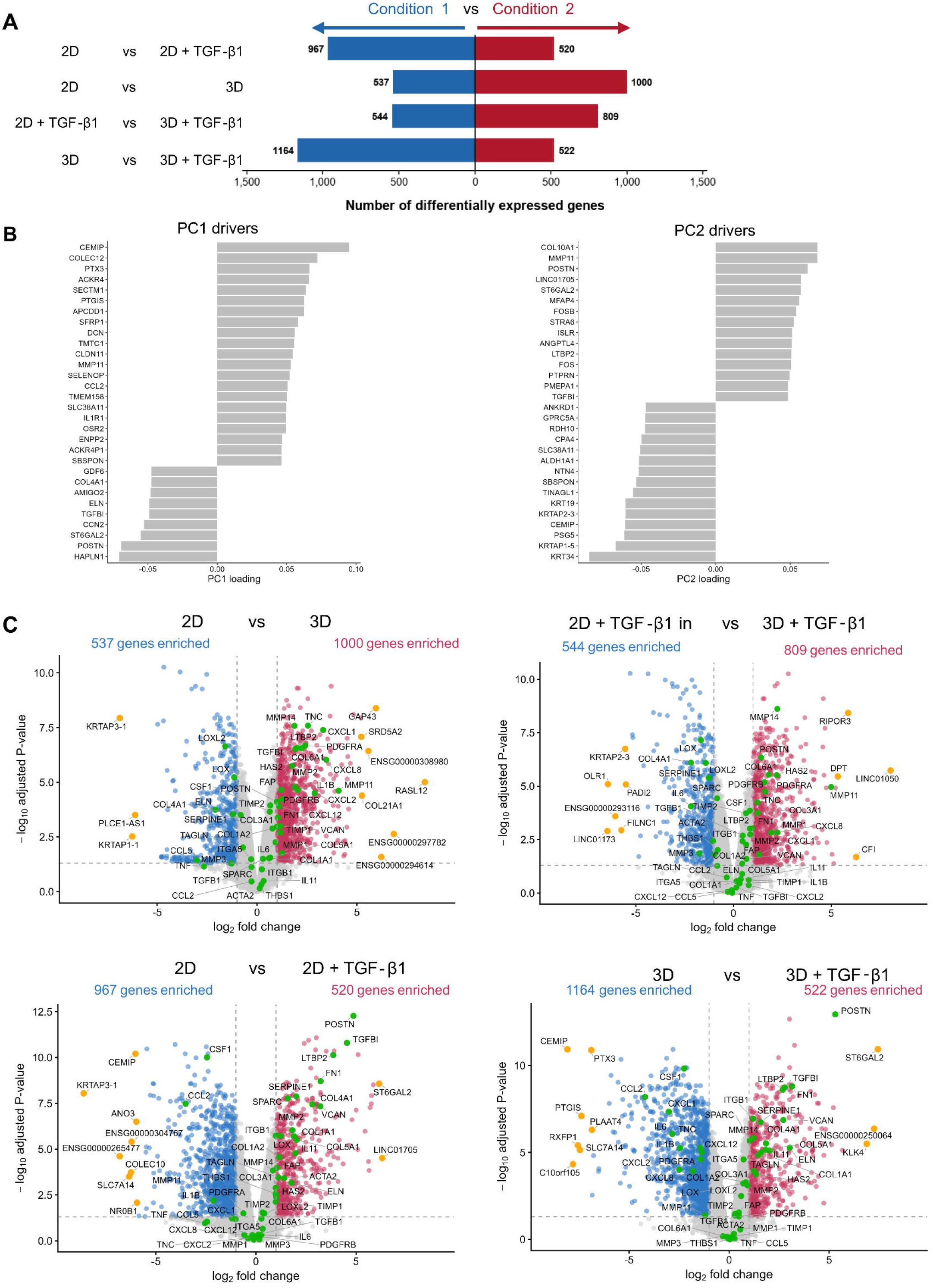
Additional RNA-seq analysis. **A)** Bar graph showing the number of DEGs identified for each indicated comparison. For each comparison, blue bars on the left indicate the number of DEGs increased in condition 1, whereas red bars on the right indicate the number of DEGs increased in condition 2. DEGs were defined using an adjusted *p*-value < 0.05 and a log₂ fold-change > 1. **B)** Gene loadings driving PC1 and PC2 separation in RNA-seq analysis. Bar graphs showing the top positive and negative loading genes. Loading values indicate the relative contribution of each gene to sample separation along the respective principal component. Positive and negative loading genes are associated with opposite ends of the PCA axis. **C)** Volcano plots showing differential gene expression between the indicated experimental conditions. Genes are plotted according to log₂ fold change and −log₁₀ adjusted p-value. Significantly upregulated and downregulated genes are color-coded, whereas non-significant genes are shown in grey. Selected genes included in the corresponding heatmap are highlighted in green, and the top 10 differentially expressed genes are highlighted in orange. Differential expression analysis was performed on RNA-sequencing data from n = 3 individual experiments.

**Supplementary Figure 2 :**
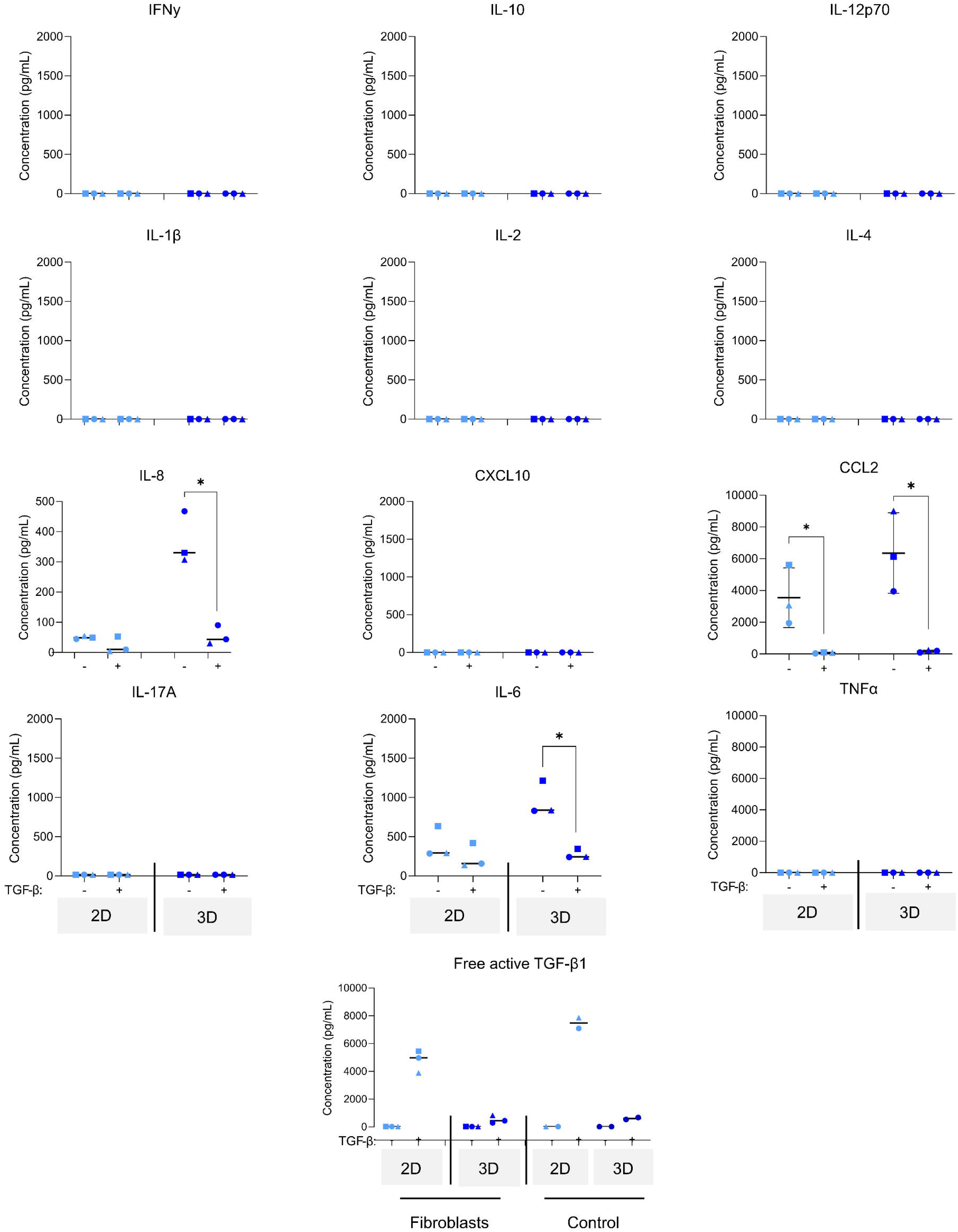
Total panel of profiled cytokine and chemokines measured in the conditioned medium of fibroblasts cultured in 2D or 3D embedded in collagen I hydrogels. The concentration of IFN-γ, IL-1, IL-12p70, IL-1β, IL-2, IL-4, IL-8, CXCL10, CCL2, IL-17A, IL-6, TNF-α, and free active TGF-β1 in medium collected from fibroblasts cultured in 2D or 3D, with and without TGF-β1. Bars show the mean ± SD of medium collected from three independent experiments. Statistical analysis was performed using paired t-test. *P < 0.05.

**Supplementary Figure 3 –.**
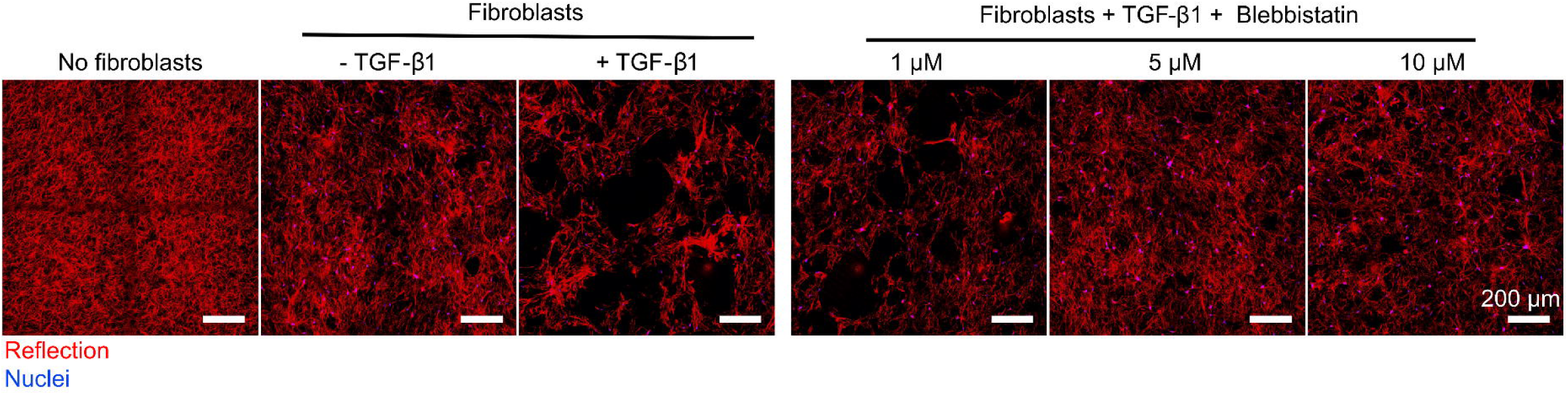
Matrix remodeling depends on acto-myosin mediated contractility. Representative maximum-intensity projections of confocal microscopy imaging showing collagen I gels with or without fibroblasts or TGF-β1-activated fibroblasts and treated with 1, 5 and 20 μM blebbistatin. Blue, Hoechst stained nuclei; red, ECM fibers detected by reflection microscopy. Scale bar = 200 μm

**Supplementary Table 1 –.** Selected gene panel used for row-scaled expression heatmap and volcano plots.

| **Functional category** | **Genes included** | **Number of genes** |
|---|---|---|
| Inflammatory mediators | CCL2, CCL5, CCL7, CCL8, CSF1, CSF2, CXCL1, CXCL2, CXCL8, CXCL10, CXCL12, IFNG, IL1B, IL1A, IL2, IL4, IL6, IL12B, IL10, IL11, IL17A, TNF | 22 |
| ECM structure | COL1A1, COL1A2, COL3A1, COL4A1, COL5A1, COL6A1, ELN, FN1, HAS2, POSTN, TNC | 11 |
| ECM remodelling | MMP1, MMP2, MMP3, MMP9, MMP11, MMP14, LOX, LOXL2, TIMP1, TIMP2 | 10 |
| Myofibroblast / TGF-β | ACTA2, CTGF, FAP, LTBP2, PDGFRA, PDGFRB, SERPINE1, TAGLN, TGFB1, TGFBI, | 10 |
| Cell–matrix interaction | ITGA5, ITGB1, THBS1, SPARC, VCAN | 5 |

